# TGFβ treatment alters the scleral fibroblast migratory response to cyclic strain in an ERK-dependent manner

**DOI:** 10.1101/2022.05.14.491950

**Authors:** Ann Mozzer, Ian Pitha

## Abstract

**Purpose:** Myofibroblasts are associated with scleral remodeling in myopia and glaucoma. We showed reduced myofibroblast alignment with topographic cues in sclera can be modeled by cyclic strain exposure of aligned myofibroblasts *in vitro*. Here, we characterize the scleral myofibroblast response to cyclic mechanical strain.

**Methods:** Human peripapillary scleral (PPS) fibroblasts were cultured on topographically aligned grooves to promote cell alignment, exposed to TGFβ (2 ng/ml) in the presence of vehicle or kinase inhibitors, and exposed to uniaxial strain (1 Hz, 5%, 12-24 hours). Alignment with grooves was determined at baseline, immediately following strain, and 24 hours after strain cessation with 0° being completely aligned and 90° being perpendicular to grooves. A wound healing assay was developed to investigate further fibroblast migration across topographic cues. Transcriptional profiling of myofibroblasts with or without strain was performed by RT-PCR and pERK, pSMAD2, and pSMAD3 levels were measured by immunoblot.

**Results:** Pre-strain alignment (6.2±1.5°) was reduced after strain (21.7±5.3°, *p*<0.0001) and restored 24 hours after cessation (9.5±2.6°). ERK, FAK, and ALK5 inhibition preserved alignment following strain; however, alignment reduction was not inhibited by ROCK, YAP, or SMAD3 inhibition. TGFβ-induced myofibroblast markers were reduced by strain. While TGFβ-induced phosphorylation of ERK and SMAD2 was unaffected by strain, SMAD3 phosphorylation was reduced (*p*=0.0004). Wound healing across grooves was enhanced by ROCK and SMAD3 inhibition but not ERK or TGFβR1 inhibition.

**Conclusions:** Strain-induced myofibroblast migration across topographic confinement is ERK dependent and associated with pSMAD3 inhibition. These results provide insight into potential mechanisms of pathologic scleral remodeling.

**Precis:** Glaucomatous scleral remodeling is driven by cellular activity. Here we find that scleral myofibroblasts have an exaggerated response to mechanical strain that is ERK dependent, associated with pSMAD3 inhibition and mitigated TGFβ signaling.

## 1. Introduction

Glaucomatous vision loss occurs because of retinal ganglion cell (RGC) injury and death due to axonal injury that is initiated at the optic nerve head (ONH). One pathway of axonal injury occurs through translation of IOP-induced mechanical stress to OHN axons and astrocytes which leads to axonal transport blockade and optic nerve astrocyte process reorganization. IOP-induced strain is translated to the ONH through the scleral wall (hoop stress) and across the translaminar pressure gradient which is determined by the pressure differential between IOP and optic nerve cerebral spinal fluid (CSF)^1^. The peripapillary sclera (PPS) is the scleral portion of the ONH that translates hoop and translaminar IOP strain to ONH axons and therefore is an area of significant relevance in glaucoma progression. In glaucomatous eyes, sclera stiffens, and collagen fibers of the PPS become less aligned^1^. These changes alter the effect of IOP level on ONH function; however, the mechanisms of sclera and PPS remodeling have not been fully explored^2, 3^. While passive and active processes could be involved in scleral remodeling, the bulk of scleral remodeling is likely accomplished though cellular activity.

The sclera is relatively paucicellular but contains populations of macrophages, stem cells, and fibroblasts^4–6^. Resident fibroblasts are the main scleral cellular component and can be found throughout the sclera with high cytoskeletal alignment to surrounding extracellular matrix (ECM)^7^. Oglesby and colleagues showed fibroblast proliferation after IOP elevation and increased aSMA expression which is a marker for myofibroblasts - a cell type involved in tissue remodeling throughout the body^8^. Additionally, increased scleral myofibroblast markers are found in myopia models which further suggests that this cell type plays a role in diseases of scleral remodeling^9^. Since scleral fibroblasts and myofibroblasts are closely associated with scleral ECM, respond to IOP elevation, and IOP-induced mechanical strain contributes to ONH dysfunction, the response of these cell types to extracellular cues present in glaucomatous sclera is highly relevant to glaucoma pathogenesis. These findings could also have relevance to lamina cribrosa (LC) remodeling as mechanosensitive lamina cribrosa cells are hypothesized to drive glaucomatous LC remodeling^10^.

Fibroblasts in glaucomatous sclera are exposed to multiple, interacting extracellular cues. IOP-induced mechanical strain is highly complex because IOP is influenced by variations in pulse pressure pulse, blinking, eye movements, and body position^11^. These influences are highly variable and cause pressure spikes and troughs on a second-to-second time scale. It is not clear whether resident scleral fibroblasts sense all IOP variation because scleral biomechanical properties could shield fibroblasts from some of these variations. Mechanical strain and glaucoma can influence scleral cytokine signaling. TGFβ levels were elevated in glaucomatous ONH^8, 12^. TGFβ is a profibrotic cytokine that drives myofibroblast differentiation and can be held in an inactive state until release from extracellular reservoirs. Lastly, scleral collagen fibrils are regionally specialized within the PPS containing an annulus of circumferentially oriented fibers that protect the optic nerve from mechanical stress^3, 13, 14^. This organization is markedly different from the basket weave pattern seen in peripheral scleral tissue. Glaucomatous eyes partially lose PPS collagen fiber alignment which could reduce the ability of the PPS to protect the optic nerve from IOP-induced mechanical strain.

The effect of topographical and mechanical cues on cellular behavior have been extensively studied. Cells recognize topographical cues ranging from sub-micrometer to ten micrometers in size and organize their cytoskeleton to align and migrate along these cues. This phenomenon is known as contact guidance and is governed by focal adhesions, cell cytoskeletal elements, and cellular contractile machinery^15^. Adhesive cells respond to uniaxial cyclic strain by exhibiting a strain avoidance response in which cells reorient to a position with their long axis almost perpendicular to the strain direction^16, 17^. Topographical and strain cues can compete in directing cellular alignment – when uniaxial strain is applied in a direction parallel to topographic cues^18^. When conflicting signals are present, the ability of strain avoidance cues to overcome contact guidance depends on the degree and duration of strain – with longer and more intense strain allowing cells to overcome contact guidance cues and reorient^17, 19^. The extent to cell type and cytokine exposure can influence the balance of these signals has not yet been investigated.

In normal human sclera, we measured a high degree of cellular cytoskeletal alignment with surrounding collagen ECM fibrils; however, alignment was reduced in aSMA-expressing myofibroblasts^7^. When examined *in vitro*, myofibroblasts oriented with contact guidance cues in the absence of strain. We therefore hypothesized that myofibroblasts more readily overcome contact guidance cues when presented with conflicting cues from mechanical strain. This hypothesis was validated by our finding that TGFβ-induced myofibroblast differentiation in combination with cyclic uniaxial strain promoted cell reorientation and decreased alignment with surrounding topographic cues. This finding was relevant to glaucomatous scleral remodeling as scleral myofibroblasts are exposed to IOP strain in glaucomatous eyes and an altered migratory phenotype could underly the observed reduction of collagen fiber alignment seen in glaucomatous eyes. Here, we explore this migratory phenomenon further by investigating the molecular signaling pathways that underlie this response.

## 2. Materials and Methods

### 2.1 Materials

Kinase inhibitors were obtained from the following suppliers: PF573228 (Cayman Chemical Company, Ann Arbor, MI), SIS3 (APExBIO, Houston, TX), SCH772984 (APExBIO), verteporfin (TOCRIS, Minneapolis, MN), galunisertib (R&D Systems, Minneapolis, MN), Y27632 (TOCRIS), and fasudil (TOCRIS). Recombinant human TGFβ1 and TGFβ2 were obtained from R&D Systems. Collagen I (rat tail) was obtained from Thermo Fisher (Waltham, MA).

### 2.1 Fibroblast culture

Methods for isolation and culture of PPS were described previously in detail.^20^ Donor eyes were obtained from the NDRI within 24 hours of enucleation. Cell lines were isolated from eyes of three European-derived persons without a clinical history of glaucoma or pathologic myopia: (1) a 76-year old Caucasian man, (2) a 97-year old Caucasian woman, and the left (3) and right (4) eyes from a 56-year old Caucasian man with no past ocular history other than cataracts. Briefly, a 2-mm wide scleral band surrounding the ONH was isolated and cut into 1 by 1 mm sections. Sclera pieces were then placed on 35 mm collagen-coated, tissue culture dishes containing RPMI-1640 medium, 20% fetal bovine serum (FBS), non-essential amino acids, 1% penicillin/streptomycin, and sodium pyruvate. After 14 days, cells were passaged and cultured in Dulbeccos’s modified Eagle’s medium (DMEM) with 10% FBS, 1% penicillin/streptomycin, and sodium pyruvate. All experiments were conducted on cells between passages 3 and 10 using passage media with 1% FBS.

### 2.2 Cell strain experiments

Cyclic cellular strain experiments were conducted using a Nanosurface Cytostretcher © (Nanosurface Biomedical Inc., Seattle, WA) and collagen-coated chambers with topography in parallel with direction of strain (Nanosurface Biomedical). Dimensions of the grooves and ridges of the parallel topography surface were 800 nm with a depth of 600 nm^21^. For orientation, immunofluorescence, and transcription experiments 1.44 cm^2^ chambers were used while 25 cm^2^ chambers were used for immunoblot experiments. After seeding, PPS fibroblasts were allowed to settle for 48-hours in media containing 1 % FBS prior to strain. Kinase inhibitors were added 2 hours prior to strain and TGFβ1 (R&D Systems, Minneapolis, MN) (2 ng/ml) was added 1 hour prior to strain.

### 2.3 Fibroblast orientation analysis

Cell body analysis was performed on light microscope images of cells taken using a 10x objective lens on a Nikon Eclipse TS100 microscope (Nikon Inc., Melville, NY) with a SPOT Insight digital camera (Diagnostic Instruments, Inc., Sterling Heights, MI). At least 12 images were taken per chamber. Orientation was determined manually for each cell using the Measure function of ImageJ for at least 240 cells per chamber by a masked grader.

For nuclei and actin analysis, cells were immediately fixed in 4% paraformaldehyde after cyclic strain, labelled with nuclear (DAPI, Thermo Fischer Scientific, Waltham, MA) and fibrillar actin (phalloidin Alexa Flour 568, Thermo Fischer Scientific, Waltham, MA) stains, and imaged using a EVOS cell imaging system and a 10x objective lens (Thermo Fischer Scientific, Waltham, MA). Images were analyzed using ImageJ using the Shape Descriptors plugin with parameters 10-100 to quantify and characterize nuclei^22^. Presence or absence of stress fibers was determined by a trained masked grader.

### 2.4 Modified wound healing assay

Fibroblasts were seeded in collagen-coated, 24-well smooth bottomed or topographically grooved polystyrene plates (Nanosurface Biomedical Inc., Seattle, WA) and allowed incubate for 24 hours. Two hours prior to wound creation, cells were treated with kinase inhibitor with and without TGF (2 ng/ml). Each well received one scratch wound created with a 200 *μ*l pipette tip. On smooth plates, a vertical wound was created. On patterned plates, two types of wounds were created (1) one perpendicular to the direction of the grooved surface (perpendicular) and (2) one parallel to the direction of the grooved surface (parallel). Wound healing was followed serially by imaging with a Nikon Eclipse TS100 microscope (2 images per well; 4 wells per condition) and images were analyzed by a masked grader using ImageJ. Would healing was calculated as relative wound area compared to wound area measured immediately after wound.

### 2.6 Transcription analysis

Following strain, cells were immediately washed with phosphate buffered saline (PBS, 4°C) and RNA isolated using Rneasy Micro Kit (Qiagen, Valencia, CA) and cDNA was generated using the High-Capacity cDNA Reverse Transcription Kit (Thermo Fisher Scientific, Waltham, MA) according to the manufacturer’s instructions. Gene expression levels were normalized to glyceraldehyde 3-phosphate dehydrogenase (GAPDH) mRNA and reported as fold change over fellow control eye. Primer sequences used are listed in **Supplemental Table 1.** The mRNA expression of genes was evaluated using a QuantiStudio 3 Real-Time PCR System (Thermo Fisher), SYBR Green Real-Time PCR Master Mix (Thermo Fisher Scientific), and the 2^−ΔΔCt^ method.

### 2.7 Immunoblot analysis

Cell lysates were resolved on 10% SDS-PAGE gels under reducing conditions. Proteins were then transferred to nitrocellulose membranes that were blocked with TBS containing 5% non-fat milk and 5% bovine serum albumin with 0.1% Tween 20 and incubated with antibody (**Supplemental Table 2**). Antigen-antibody reactions were detected using chemiluminescence and bands were quantified by Image Lab ^™^ Software (Biorad, Hercules, CA).

### 2.7 Statistical analysis

All values are mean ± standard deviation (SD). Unless otherwise noted, means were compared using one-way analysis of variance test (ANOVA) with Dunnett post-hoc tests to correct for multiple comparisons. Data normality (Gaussian distribution) was assessed by Shapiro-Wilk normality and D’Agostino-Pearson tests.

## 3. Results

### 3.1 Cyclic cell strain reduced contact guidance in TGFβ treated cells

Fibroblasts cultured on a surface with grooved topography align with the long axis of the grooves. We previously demonstrated that the combination of TGFβ treatment and cyclic uniaxial strain applied parallel to the direction of these grooves reduced fibroblast cell body alignment with surrounding topography ^7^. Here, we expand on those findings by examining fibroblast-ECM alignment at baseline, following 18 hours of cyclic strain (1 Hz, 5% strain), and after a 24-hour recovery period during which the strain was stopped. Fibroblasts placed under strain for 18 hours without TGFβ-treatment maintained alignment with topographic cues despite strain-induced cues to migrate to a perpendicular orientation. Pretreatment with TGFβ which induces myofibroblast differentiation; however, caused a significant loss of cell body-groove alignment after cell strain. Interestingly, there was a bimodal distribution with two peaks – one population of cells retained alignment and the other cell population had reduced alignment that peaks at a misalignment of approximately 20-30°. After a 24-hour recovery period, alignment was restored to baseline levels (**Figure 1**).

**Figure 1.**
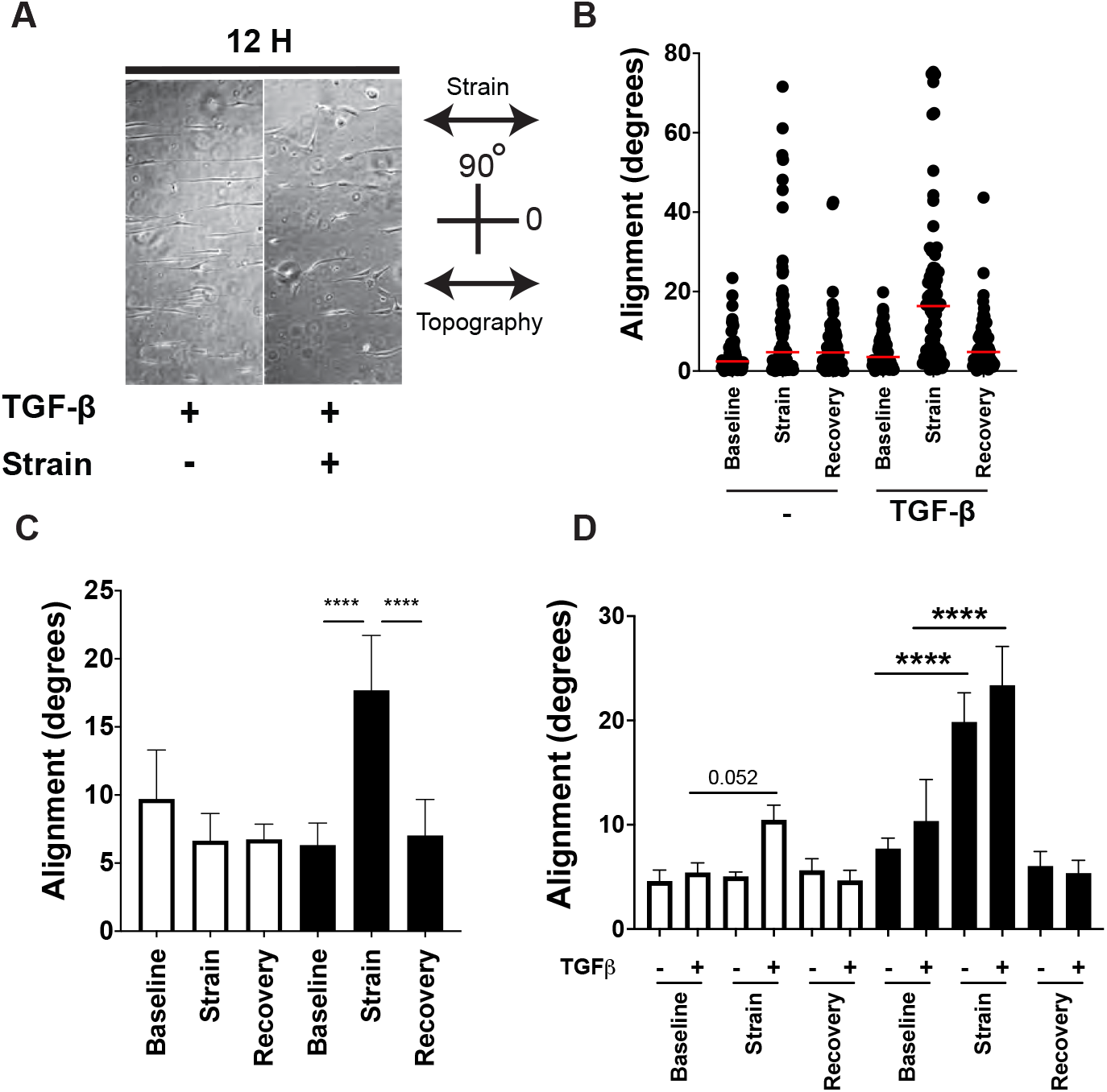
Reversible loss of topographic alignment after strain of TGFβ-treated cells. (A) Light microscope image (10x magnification) of TGFβ-treated PPS cells prior to and immediately after cyclic strain. (B) Cell alignment from a single experiment showing a bimodal distribution of alignment loss after strain of TGFβ-treated PPS cells. (C) Alignment loss across multiple experiments (*n*=3) (white bars = no TGFβ; black bars = TGFβ treatment). (D) Alignment loss after exposure to low (3%, white bars) and high (15%, black bars) strain. **** *P*<0.0001

We next tested whether this effect was dependent on the intensity of mechanical strain by performing the same study at low (3%, 1 Hz) and high (15%, 1 Hz) strain intensities. When placed under low strain for 18 hours, there was a non-significant trend for cell body reorientation following TGFβ treatment but not under TGFβ-free conditions. Under high strain conditions, however, both TGFβ treated and TGFβ-free conditions changed cellular orientation. These findings suggest that TGFβ treatment resets the rheostat for cell reorientation to occur at lower strain intensities and that this effect does not require TGFβ-treatment under high strain conditions.

### 3.2 Nuclear aspect ratio and orientation is altered by cell strain

Cytoskeletal components link the outer cell membrane and the nucleus^23^. Through this linkage and activation of signaling molecules such as YAP, ERK, and rho-kinases, external mechanical signals can initiate alterations in nuclear shape and cellular position^24^. Cell migratory activity alters nuclear characteristics as nuclear aspect ratio changes during different phases of cell migration – becoming more rounded as cells translate from persistent to hesitant migration phases ^25^. Due to this known connection between external signaling, cell migration, and nuclear dynamics, we hypothesized that strain-induced alterations in cell alignment would be paralleled by changes in nuclear shape and alignment. To test this hypothesis, we next examined changes in nuclear size, aspect ratio, and alignment in the presence of cyclic strain and TGFβ treatment. Cells were exposed a uniaxial strain as described in the previous section and immediately after completing 18 hours of cyclic strain, cells were fixed, and nuclei were stained using DAPI. This approach allowed a non-biased analysis of aspect ratio, size, and orientation using ImageJ. When exposed to cyclic strain and TGFβ, nuclear aspect ratio decreased and alignment with surrounding topographic cues was reduced when compared to control conditions (**Figure 2**). There was also a non-significant trend of nuclear enlargement.

**Figure 2.**
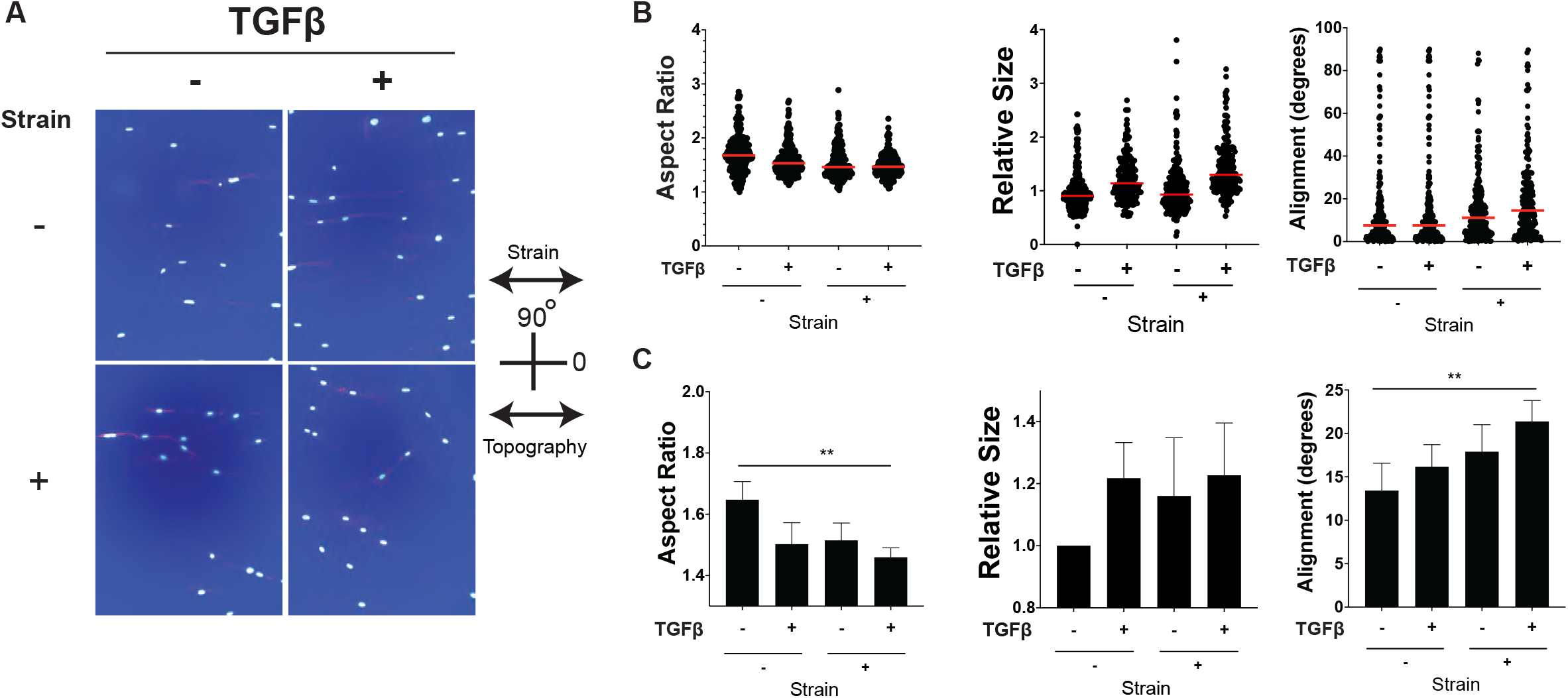
Alterations in nuclear aspect ratio and alignment. (A) Microscopic images of DAPI stained nuclei (10x magnification) of PPS cells allowed to align on topographic grooves and then exposed to TGFβ and cyclic strain (5%, 1 Hz). Nuclear characteristics from a single experiment (B) and across multiple experiments (C)(*n*=3). ** *P*<0.01

### 3.3 ERK1/2 is necessary for reduction of cell body alignment

TGFβ signals through canonical (SMAD2/3) and non-canonical (ROCK, ERK, p38, JNK, YAP) pathways to induce myofibroblast differentiation in fibroblasts^26^. Each of these signaling pathways contribute to the global TGFβ response, however, selective pathway inhibition can contextualize the cellular response and modify cellular behavior^27^. We therefore hypothesized that migratory reorientation with strain could be selectively regulated by specific TGFβ signaling pathways. To test the role of canonical and non-canonical TGFβ signaling in regulation of cell alignment, we pretreated aligned cells with inhibitors of TGFβ signaling immediately prior to TGFβ treatment and cyclic strain exposure. Alignment with extracellular topography was then determined at baseline, immediately after 18 hours of cyclic strain, and after a 24-hour recovery period (**Figure 3**). Inhibition of the TGFβ receptor type I (Alk5) with galunisertib (10 *μ*M) prevented loss of alignment following cyclic strain; however, SMAD3 inhibition with SIS3 (5 *μ*M) or YAP inhibition with verteporfin (3 *μ*M) did not prevent loss of alignment nor recovery after strain cessation. ROCK inhibition with Y27632 (10 *μ*M) or fasudil (2 *μ*M) did not prevent loss of cell-ECM alignment but did prevent recovery after strain cessation. ERK1/2 and focal adhesion kinase (FAK) inhibition with SCH772984 (500 nM) and PF573228 (5*μ*M), respectively, did, however, prevent cell strain-induced loss of alignment. Together, these findings demonstrate that while markers of myofibroblast differentiation can be reduced by different kinase inhibitors only inhibition of ERK1/2, FAK, or Alk5 prevented loss of cell alignment.

**Figure 3.**
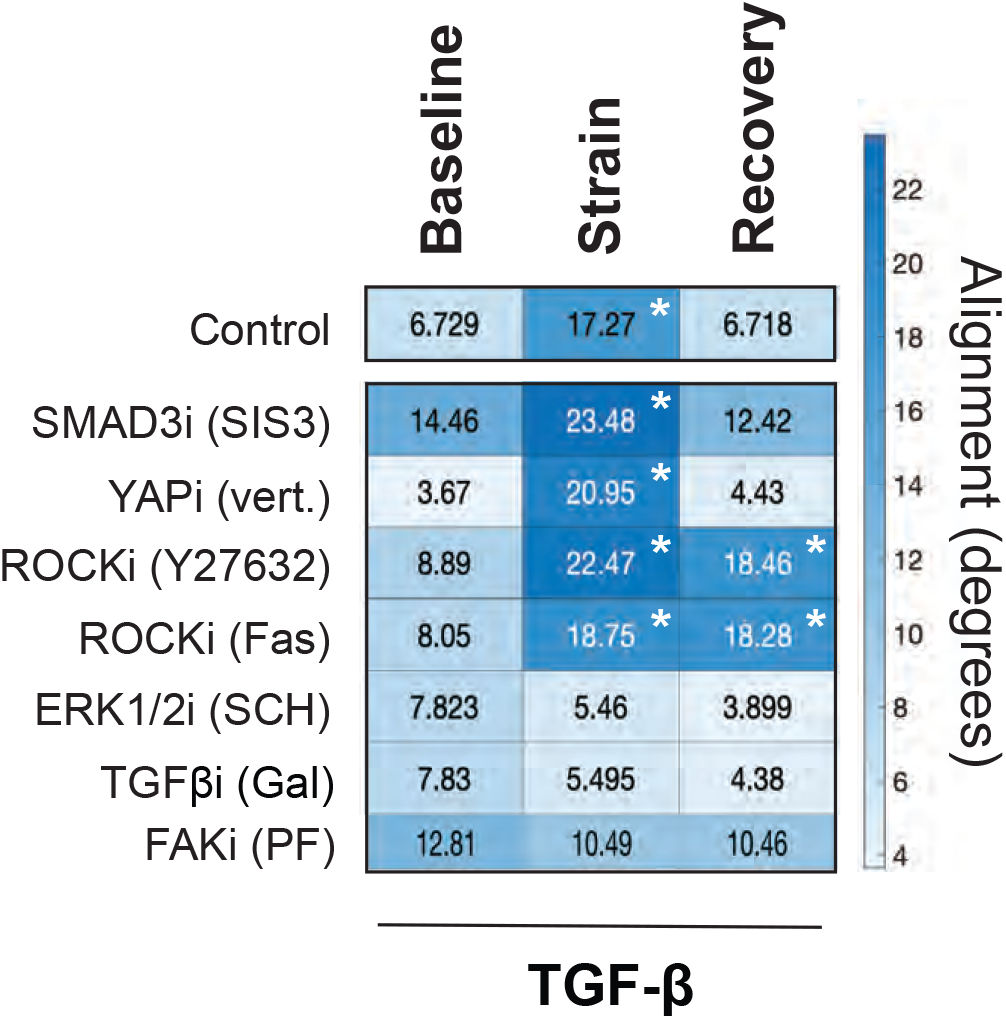
Reduction of strain-induced alignment loss by ERK, TGFβ, and FAK inhibitors. Heatmap of alignment with topographic grooves at baseline, immediately after strain (5%, 1 Hz, 18 hours), and following a 24-hour recovery period. Experiments were performed with vehicle (control) and in the presence of kinase inhibitors (*n*=2). * *P*<0.05 compared to baseline measurement.

### 3.4 ERK-inhibition reduces cell migration across topographic barriers

These results suggested that cyclic strain modifies myofibroblast migratory behavior by selectively targeting specific signaling pathways downstream from TGFβ. An unresolved question regarding this phenomenon remained whether fibroblast or myofibroblast migration across topographic barriers could be induced in the absence of mechanical strain. To address this question, we developed a modified wound healing assay to test the extent to which these downstream pathways alter contact guidance in the absence of cell strain. In the assay, fibroblasts were seeded on smooth or grooved culture surfaces. A scratch was then created on the smooth surface topography and one of two scratches was created on the grooved surfaces: (1) perpendicular to the surface grooves or (2) parallel to surface grooves (**Figure 4**). Cell migration across the wounded area was then followed. Under baseline conditions wound healing across a smooth surface or perpendicular scratch was significantly faster than healing across parallel wounds (**Figure 4**). TGFβ1 and TGFβ2 treatment accelerated wound healing across smooth and perpendicularly oriented wounds but not across parallel oriented wounds (**Figure 4**).

**Figure 4.**
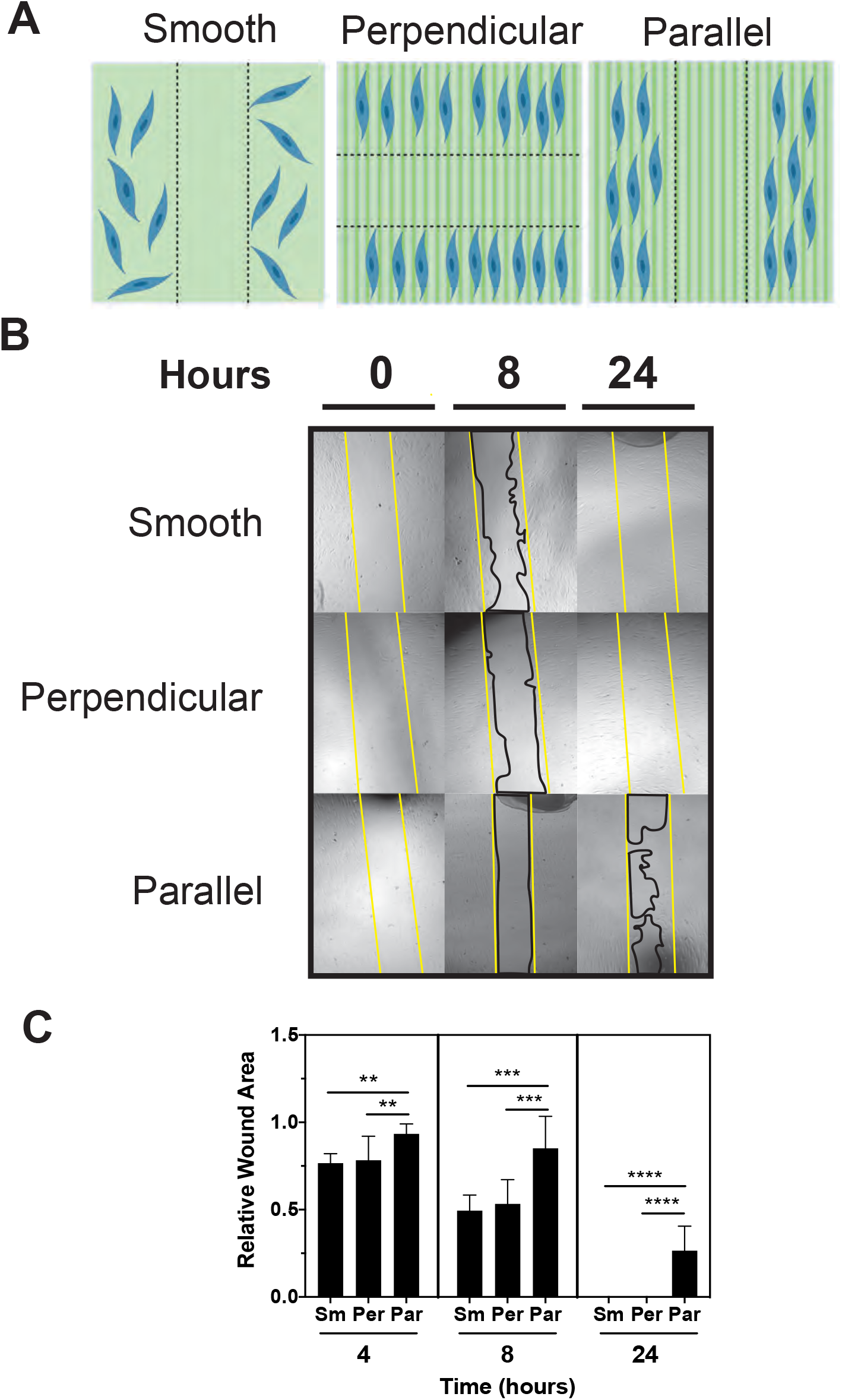
Modified wound healing assay. (A) Illustration of wounds created on smooth (Smooth) and topographically grooved (Parallel and Perpendicular) plates at time zero. (B) Light microscopy images of wounds at baseline (0), 8 hours, and 24 hours after wound creation. Yellow lines demarcate the border of the initial wound and black lines demarcate cell migration. (C) Relative wound area across smooth (Sm), perpendicular (Per), and parallel (Par) wounds (*n*=4). ** *P*<0.01, *** *P*<0.001, **** *P*<0.0001

Next, we tested migration across these wounds after treatment with kinase inhibitors and found responses varied depending on the kinase inhibitor used and the presence of TGFβ1 (**Figure 5**). SMAD3 inhibition reduced healing on smooth surface in the absence of TGFβ1 treatment but enhanced healing across parallel oriented wounds in the presence of TGFβ1. ROCK inhibition enhanced healing across smooth and parallel but not perpendicular wounds. In the presence of TGFβ, ROCK inhibition only enhanced healing across parallel oriented wounds. In contrast to these results, ERK1/2 inhibition reduced healing across smooth and perpendicular oriented wounds but not parallel wounds. In the presence of TGFβ1, ERK inhibition inhibited healing across perpendicularly oriented wounds, but this effect was only manifest at 24 hours after wounding. These findings highlighted that ERK1/2 signaling plays a necessary role in multiple cell migratory phenotypes while SMAD3 and ROCK inhibition enhanced cell migration across topographic barriers (across the parallel wounds). Enhanced migration after SMAD3 inhibition, however, was only detected in the presence of TGFβ1 treatment.

**Figure 5.**
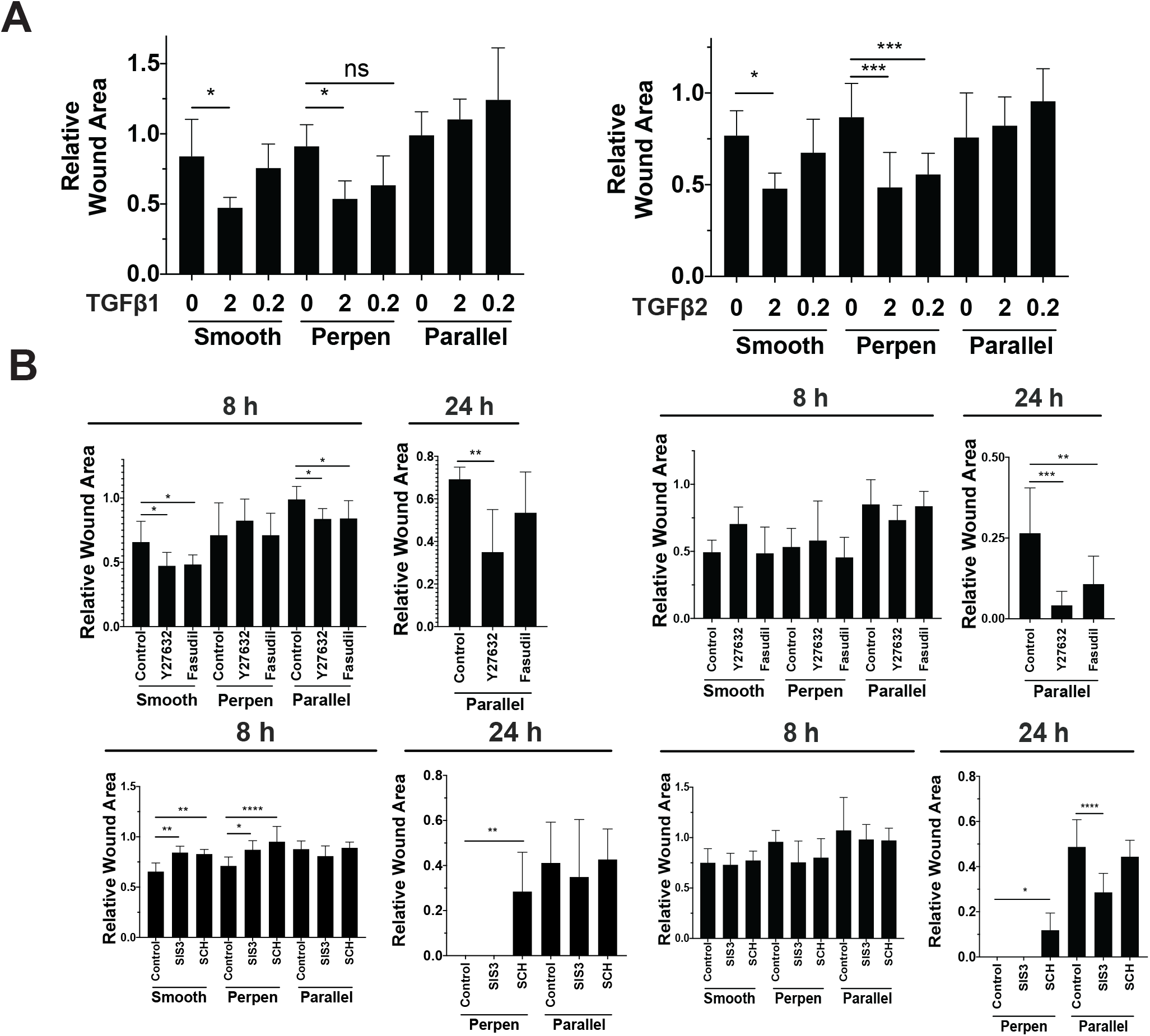
Kinase inhibitors alter PPS fibroblast migration across topographic grooves. (A) Wound healing following treatment with TGFβ1 and TGFβ2 (0, 2, and 0.2 ng/ml). (B) Wound healing without (left) and with (right) TGFβ1 treatment (2 ng/ml) after kinase inhibitor treatment (*n*=4).

### 3.5 Cyclic strain mitigates TGFβ-induced transcription of myofibroblast markers

Cyclic strain can induce myofibroblast differentiation in PPS cells^28^. This effect was dependent on strain conditions (requiring greater strain and more rapid frequency) and varied between primary cell lines tested with most lines undergoing myofibroblast differentiation but select lines undergoing a reduction in myofibroblast markers. We extended on these previous studies by testing the influence of cyclic strain on the cellular response to TGFβ-treatment. We found TGFβ-induced transcription of myofibroblast markers was mitigated by cyclic strain and that this result was consistent across cell lines isolated from different donor eyes (**Figure 6**). For example, smooth muscle actin (aSMA) transcription was induced 43.5±8.5 fold by TGFβ treatment in the absence of strain but only 23.7±9.2 fold when treated with TGFβ and exposed to cyclic strain (*p*=0.01). A similar mitigation of the transcriptional response to TGFβ treatment upon cyclic strain was seen with additional markers of myofibroblast differentiation, including collagen (COL1A1), myosin light chain kinase (MYLK1), asporin (ASPN), thrombospondin 1 (TSB1), and periostin (POSTN). When immunoblot analysis was performed, TGFβ-induced levels of myofibroblast markers aSMA and pro-collagen type I were similarly reduced by cyclic strain. Cells were then fixed immediately after stretch and nuclei and actin fibrils were imaged. Cyclic strain with TGFβ treatment reduced stress fiber formation versus TGFβ alone and led to a diffuse rather than fibrillar actin staining pattern (**Figure 6**). Taken together these results show that cyclic strain mitigates but does not completely inhibit the fibroblast response to TGFβ treatment.

**Figure 6.**
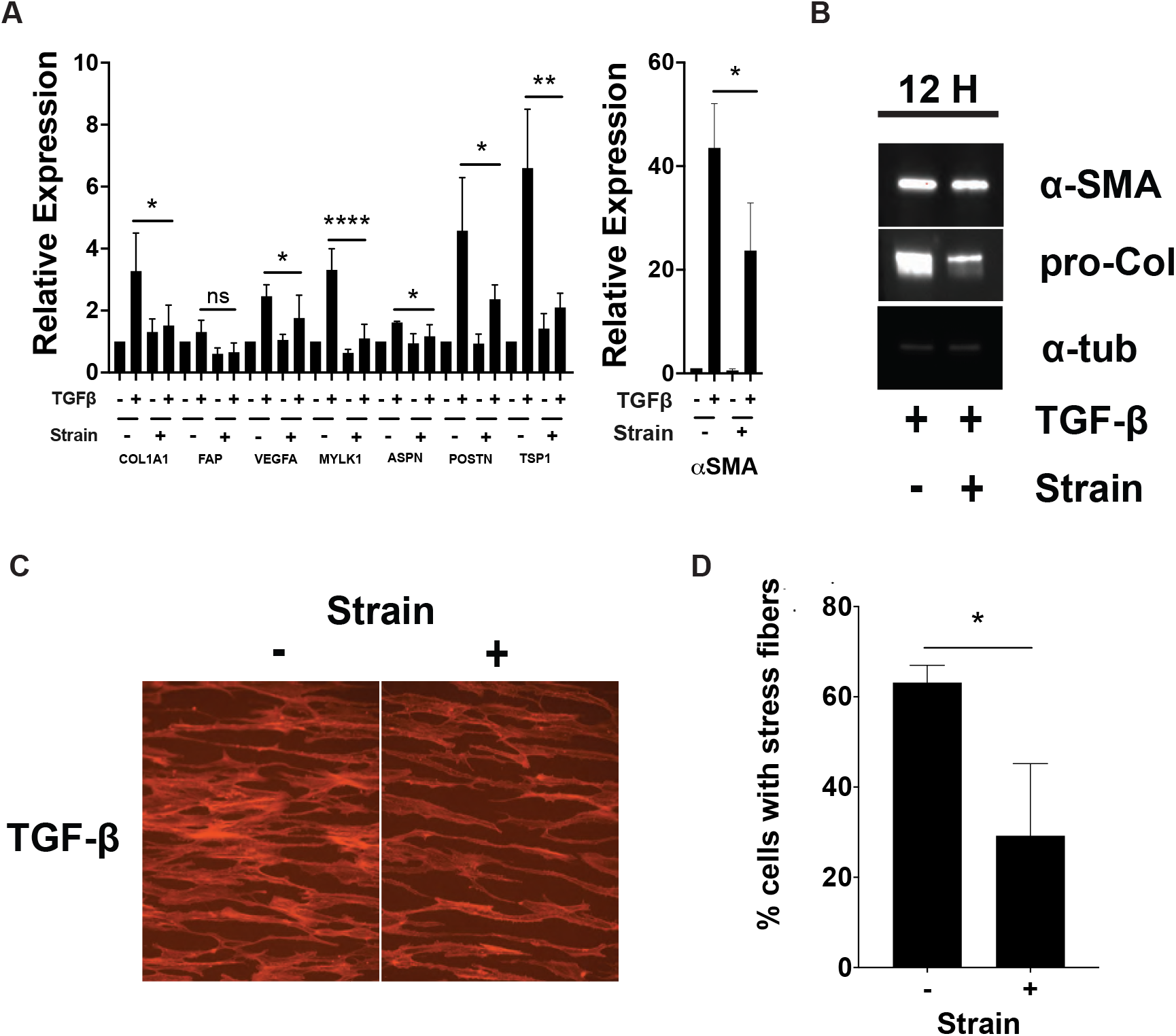
Cyclic strain mitigates TGFβ-induced effects. (A) Transcription of myofibroblast markers with and without TGFβ treatment and cyclic strain as measured by RT-PCR (*n*=4). (B) Immunoblot analysis of aSMA and pro-collagen expression in the lysate of TGFβ-treated PPS cells with and without strain. (C) Immunofluorescence images of fibrillar actin (red=phalloidin) in PPS cells exposed to TGFβ and cyclic strain. (D) Quantification of % cells with stress fibers in treated PPS cells (*n*=3).

We hypothesized that the mitigated transcriptional response was due to a reduction in the activity of downstream TGFβ signaling molecules. After TGFβ treatment, cyclic strain did not affect TGFβ-induced SMAD2 and ERK1/2 phosphorylation. Levels of pSMAD3, however, were reduced in TGFβ-treated cells exposed to cyclic strain compared to cells only treated with TGFβ (**Figure 7**). These results suggest that cyclic strain reduces the transcriptional response to TGFβ treatment by specifically targeting SMAD3 phosphorylation. In targeting SMAD3 phosphorylation but not SMAD2 or ERK1/2 phosphorylation, cyclic strain then permits a non-aligned cell body and nuclear phenotype.

**Figure 7.**
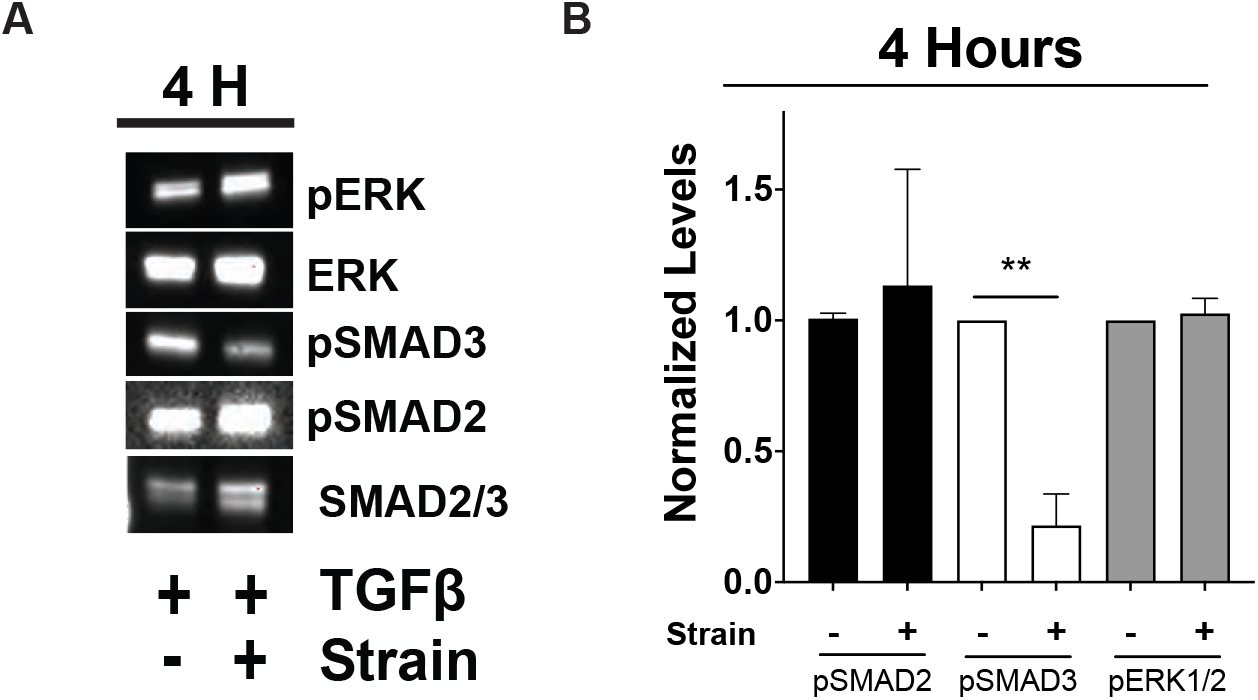
Cyclic strain selectively reduces TGFβ-induced pSMAD3. (A) Immunoblot of lysates from PPS cells taken 4 hours after TGFβ (2 ng/ml) and cyclic strain (5%, 1Hz) exposure. (B) Densitometric analysis of phosphorylated protein levels relative to ERK and SMAD2/3 levels (*n*=3).

## 4. Discussion

Here, we describe four main findings regarding the fibroblast and myofibroblast response to cyclic mechanical strain. First, TGFβ treatment reversibly alters the fibroblast response to cyclic strain by allowing cell body reorientation in the presence of contradictory topographic cues. Second, TGFβ-mediated cell reorientation is ERK-dependent and does not require SMAD3 activity. Third, cyclic strain mitigates the fibroblast transcriptional response to TGFβ treatment and reduces markers of myofibroblast differentiation. Lastly, cyclic strain selectively reduced pSMAD3 while leaving pSMAD2 and pERK levels unaffected.

Myofibroblasts had an altered and reversible response to mechanical strain. Fibroblasts and myofibroblasts exhibited contact guidance and aligned along the long axis of grooved topography under baseline conditions, but only myofibroblasts lost alignment after 3 and 5% cell strain. This finding has implications to sclera remodeling as (1) scleral myofibroblasts are enriched in glaucoma and myopia models ^8, 9^ and (2) altered collagen fibril alignment (loss of anisotropy) is found in glaucomatous PPS ^3, 29, 30^ and in posterior scleral in high myopia ^31^. We and others have hypothesized that pathologic sclera remodeling that occurs in these conditions is driven by myofibroblast activity, and the results described here suggest that myofibroblast response to mechanical strain could influence their migratory and remodeling behavior. Therefore, it is possible that scleral remodeling not only requires the presence of increased myofibroblasts but also mechanical cues mediated by IOP-mediated strain that modulate myofibroblast activity. This phenomenon would have important therapeutic implications as the therapeutic goal would transition from prevention of myofibroblast differentiation to modulation of myofibroblast activity by targeting specific signaling pathways or by managing mechanical signals to prevent formation of a pathologic myofibroblast phenotype.

Fibroblast heterogeneity can be affected by regional and temporal mechanical tissue properties. Single cell RNAseq studies have documented fibroblast heterogeneity in multiple fibrotic diseases and have shown that fibroblast populations extend beyond the simple classification of inactive fibroblasts and remodeling myofibroblasts^32–35^. While there are multiple sources of fibroblast heterogeneity including embryonic origin, cytokine exposure, position within a healing wound, and cell-cell interactions an additional source of fibroblast heterogeneity is found in the close association of fibroblasts with their surrounding ECM^36–38^. Here, we show that cyclic mechanical strain modifies the myofibroblast phenotype by altering nuclear structure, decreasing cellular contact guidance, and mitigating expression of myofibroblast markers and stress fiber formation. ECM features such as stiffness, static strain, and cyclic strain increase myofibroblast markers^34, 39–41^. Strain conditions leading to myofibroblast differentiation depended on the amount and frequency of strain exposure as well as cell type with greater, “pathologic” levels of strain being more likely to induce myofibroblast differentiation^28, 42^. The finding that cyclic strain can mitigate TGFβ-signaling was described previously in dermal fibroblasts, lung fibroblasts, stem cells, and cardiomyocytes^43–45^. Additionally, reduced aSMA expression associated with FGF-induced myofibroblast dedifferentiation was shown to be ERK-dependent^46^. In agreement with these previous studies, we found mitigation of myofibroblast markers with cyclic strain and extended these findings by describing an ERK- and FAK-dependent migratory phenotype in strain-exposed cells.

Strain-induced cell reorientation was regulated by select TGFβ signaling pathways. TGFβ receptor associate kinase ALK5 inhibition reduced cell reorientation as did inhibition of ERK and FAK. ERK’s role in cell migration, nuclear structure, and cellular response to mechanical strain is well documented^47^. FAK is a gatekeeper through which extracellular mechanical signals are communicated to the intracellular space and FAK inhibition was shown previously to reduce cell migration ^48^. In contrast, YAP, ROCK, and SMAD3 inhibition did not alter fibroblast reorientation despite being delivered at dosages sufficient to reduce myofibroblast markers. These results support further a model in which myofibroblast behavior can be tuned by targeting specific signaling events either by pharmacologic treatment or by modification of the mechanical cellular environment. In this model, it is not sufficient to identify myofibroblasts by only using traditional markers, but one must also consider cellular morphology, migratory phenotype and perhaps the relative expression levels of these markers to properly characterize cells.

We found cyclic strain selectively reduced TGFβ-induced pSMAD3 levels but did not affect pSMAD2 or pERK levels. This result suggests that pSMAD3 inhibition plays an important role in permitting a migratory fibroblast phenotype and is consistent with the findings that pSMAD3 inhibition did not prevent cell alignment loss after cyclic strain, however, pSMAD3 inhibition enhanced migration across parallel scratch wounds. Despite a high level of homology between SMAD2 and SMAD3, there is evidence for unique roles for SMAD3 in fibroblast behavior. SMAD3 deletion reduced dermal wound healing time and decreased inflammation^49^.

SMAD3 but not SMAD2 deletion in fibroblasts of infarcted hearts increased the incidence of catastrophic late rupture and adverse tissue remodeling resulting in a disorganized scar^50^. Additionally, it was hypothesized that SMAD3 drives wound consolidation and collagen alignment by regulating integrin signaling, increasing focal adhesions, and encouraging aSMA recruitment to stress fibers ^51^. Selective reduction of pSMAD3 could therefore reduce stress fiber formation and permit cell migration across topographic barrier. The extent to which SMAD3 activity is relevant to the *in vivo* response of scleral fibroblasts to glaucoma and to glaucoma progression is a topic of current study.

The present study had some recognized limitations. First, strain experienced by sclera fibroblasts *in vivo* is complex and highly variable and is not fully modeled by our *in vitro* model. Second, while the grooved topography of our *in vitro* system aligned cells to better model fibroblast alignment *in vivo*, again our *in vitro* system did not fully model the complex scleral interstitium. Third, we only included cell lines isolated from non-glaucomatous eyes. In the future it would be interesting to include cell lines isolated from glaucomatous eyes, cell lines isolated from different scleral regions, and cell lines isolated from a more diverse donor pool.

---

**Supplemental Table 1.**
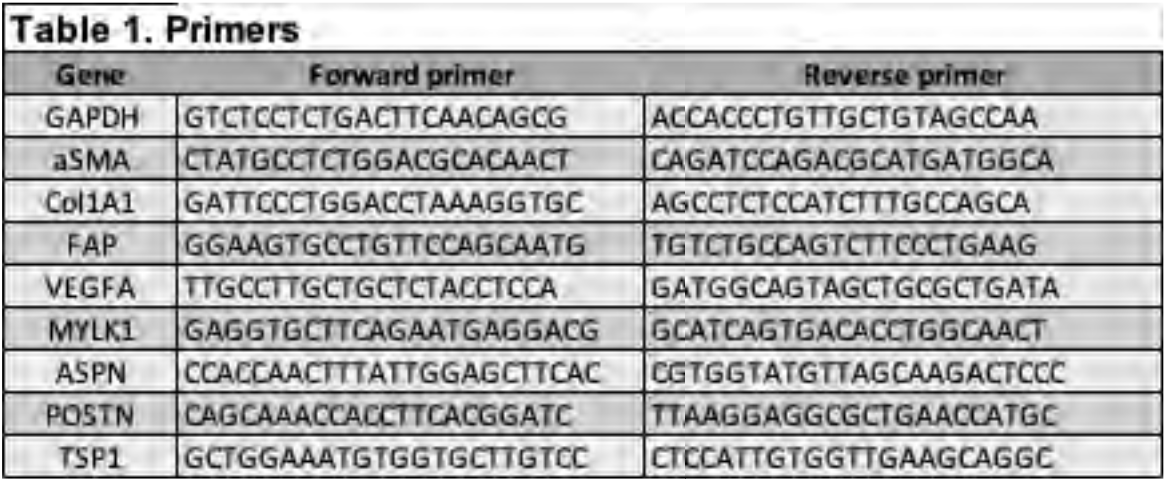

**Supplemental Table 2.**

| Table 2. Primary antibodies and stains |  |  |  |
| --- | --- | --- | --- |
| Antibody / Stain | Dilution | Species | Company |
| αSMA | 1:2000 | Ms | Sigma |
| α tubulin | 1:1000 | Ms | Abcam |
| pSMAD2 | 1:500 | Rb | Cell Signaling Tech |
| pSMAD3 | 1:500 | Rb | Cell Signaling Tech |
| SMAD2/3 | 1:1000 | Rb | Cell Signaling Tech |
| ERK | 1:1000 | Rb | Cell Signaling Tech |
| pERK | 1:1000 | Rb | Cell Signaling Tech |
| pro-collagen, type I | 1:1000 | Ms | DSHB |

## Funding/Acknowledgements

IP was supported by a RPB Career Development Award and a BrightFocus Grant. The funders had no role in study design, data collection and analysis, decision to publish, or preparation

